# A non-Hebbian code for episodic memory

**DOI:** 10.1101/2024.02.28.582531

**Authors:** Rich Pang, Stefano Recanatesi

## Abstract

Hebbian plasticity has long dominated neurobiological models of memory formation. Yet plasticity rules operating on one-shot episodic memory timescales rarely depend on both pre- and postsynaptic spiking, challenging Hebbian theory in this crucial regime. To address this, we present an episodic memory model governed by a simple non-Hebbian rule depending only on presynaptic activity. We show that this rule, capitalizing on high-dimensional neural activity with restricted transitions, naturally stores episodes as paths through complex state spaces like those underlying a world model. The resulting memory traces, which we term path vectors, are highly expressive and decodable with an odor-tracking algorithm. We show that path vectors are robust alternatives to Hebbian traces when created via spiking and support diverse one-shot sequential and associative recall tasks, and policy learning. Thus, non-Hebbian plasticity is sufficient for flexible memory and learning, and well-suited to encode episodes and policies as paths through a world model.

---

It has long been hypothesized that Hebbian plasticity— the strengthening of synaptic connections between repeatedly co-active neurons—is the neural substrate for memory formation (1–8). The canonical mechanism thought to implement Hebbian plasticity *in vivo* is spike-timing dependent plasticity (STDP), in which a synapse between two neurons potentiates if the postsynaptic neuron spikes (emits an action potential) immediately after the presynaptic neuron spikes (5, 9–11). Yet while STDP represents a compelling, intuitive mechanism for physically binding sequential neural representations together in memory, it remains unclear whether it can support a fundamental memory task: the one-shot memorization of novel complex episodic experiences (e.g. a conversation with a friend or a new route to a food source). This task is critical for rapid learning and flexible behavior, and disrupted in disorders like Alzheimer’s Disease (12).

Specifically, STDP appears ill-suited for storing information experienced only once during a realistic behavioral episode lasting seconds to minutes. First, STDP typically requires many repetitions of the coincident neural activity patterns to have an effect (10, 13–15); thus, it may act too gradually to store an episode experienced only once. Second, STDP coincidence windows are usually on the order of 10-20 ms (10)—or at best up to 100-200 ms (16)—whereas realistic behavioral intervals may be much longer (e.g. a 10 s pause between two actions; although see (17)). Such precision timing also challenges the robustness of STDP to spiking variability (18) and the ability to store memories of low-firing-rate activity patterns, since these will produce fewer coincidence events. Thus, while STDP may support multi-trial learning of tasks like pattern detection and sequence generation (5, 19, 20), the above suggest that one-shot episodic memory formation may occur via other mechanisms (Fig. 1).

**Fig. 1.**
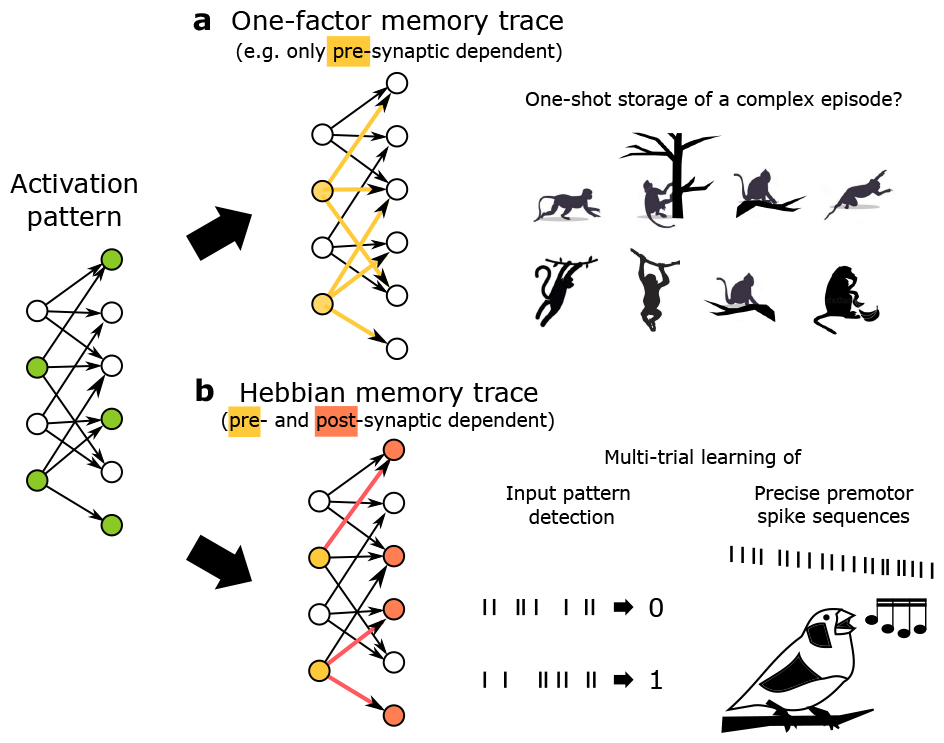
Can complex episodes with novel sequential or associative information be stored with a one-factor plasticity rule? a) Left: schematic of a basic non-Hebbian plasticity mechanism, in which synaptic potentiation is triggered only by presynaptic activity. Right: this learning rule allows for one-shot storing of complex episodes. b) Left: schematic of a memory trace stored via Hebbian plasticity, in which the synapses connecting co-active neurons strengthen. Right: After many repetitions, Hebbian STDP can learn to perform temporal pattern detection (19) or generate pre-motor spike sequences (20).

Empirically, the strongest plasticity rules thought to act in natural settings over one-shot memory and learning timescales largely do not depend on both pre- and postsynaptic spiking. One example is in hippocampal mossy fiber (MF) synapses from dentate gyrus to area CA3, which exhibit strong potentiation over multiple timescales that is thought to be triggered predominantly by presynaptic activity (21–23). For instance, it was found that naturalistic presynaptic activity triggered strong potentiation of MF synapses, via vesicle accumulation, lasting up to minutes (22). A related non-Hebbian rule is behavioral timescale synaptic plasticity (BTSP) of synapses from area CA3 to CA1 (24, 25). There, presynaptic activity is thought to create a synaptic eligibility trace that can be strongly potentiated conditional not on post-synaptic spiking but on a dendritic plateau potential occurring within a multi-second window around the presynaptic input. A third example is plasticity of synapses from Kenyon cells onto mushroom body output neurons in flies, which strongly, rapidly depress in response to presynaptic activity paired with a low-dimensional dopamine signal, rather than postsynaptic spiking (26, 27). A final pure one-factor example is activity-dependent excitability changes, which depend on a neuron’s own spiking and are also often faster than STDP (28–31).

These forms of plasticity are rapid, strong, and enduring, making them well posed for episodic memory, but they lack the intuitive local associativity of Hebbian plasticity. This raises a fundamental theoretical question: how could such rules retain in memory an intricate behavioral episode after a single experience? Could they store complex one-shot episodes containing rich sequential or associative information in a bioplausible, decodable memory trace (Fig. 1)?

In this study, we demonstrate that this issue can be resolved using a “one-factor” plasticity rule—for example, which depends only on presynaptic spiking—given an adequate systems context in which the rule is embedded. We show that the encompassing system needs three essential elements: (1) a complex state space like that underlying a world model; (2) a simple random encoding network; and (3) a decoding algorithm akin to odor-tracking or path-following—which are thought to be accessible in many animal species. We show that in this system a one-factor rule is naturally poised to store episodes as complex, information-rich paths through the state space and in a format (which we term a high-dimensional *path vector*) that enables rapid reconstruction of the episode by retracing the path with the decoding algorithm. We demonstrate the ability of this system to accomplish a range of sequential and associative one-shot memory tasks, in addition to supporting a simple but effective policy-learning scheme. In our Discussion, we outline potential improvements that could be achieved through additional gating or modulatory inputs inspired by BTSP or mushroom body plasticity (24, 26, 27). To our knowledge this is the first work showing how non-Hebbian plasticity rules could support a robust, general one-shot episodic memory system, revealing a new theoretical link between cellular physiology and the macroscopic organization of memory.

## Results

### One-factor plasticity supports storage and recall of paths through a complex state space like that of a world model

The fundamental question we address here is how a single sequentially structured experience, or episode, could be stored by a non-Hebbian one-factor rule. Our success criterion is the efficient and accurate reconstruction of the episode from the one-factor trace using biologically realistic processes. We describe the key model components—the plasticity rule, the state space, the encoding network, and the decoding algorithm (Figs. 2a to 2d)—then show how they are combined to store and retrieve an episode.

**Fig. 2.**
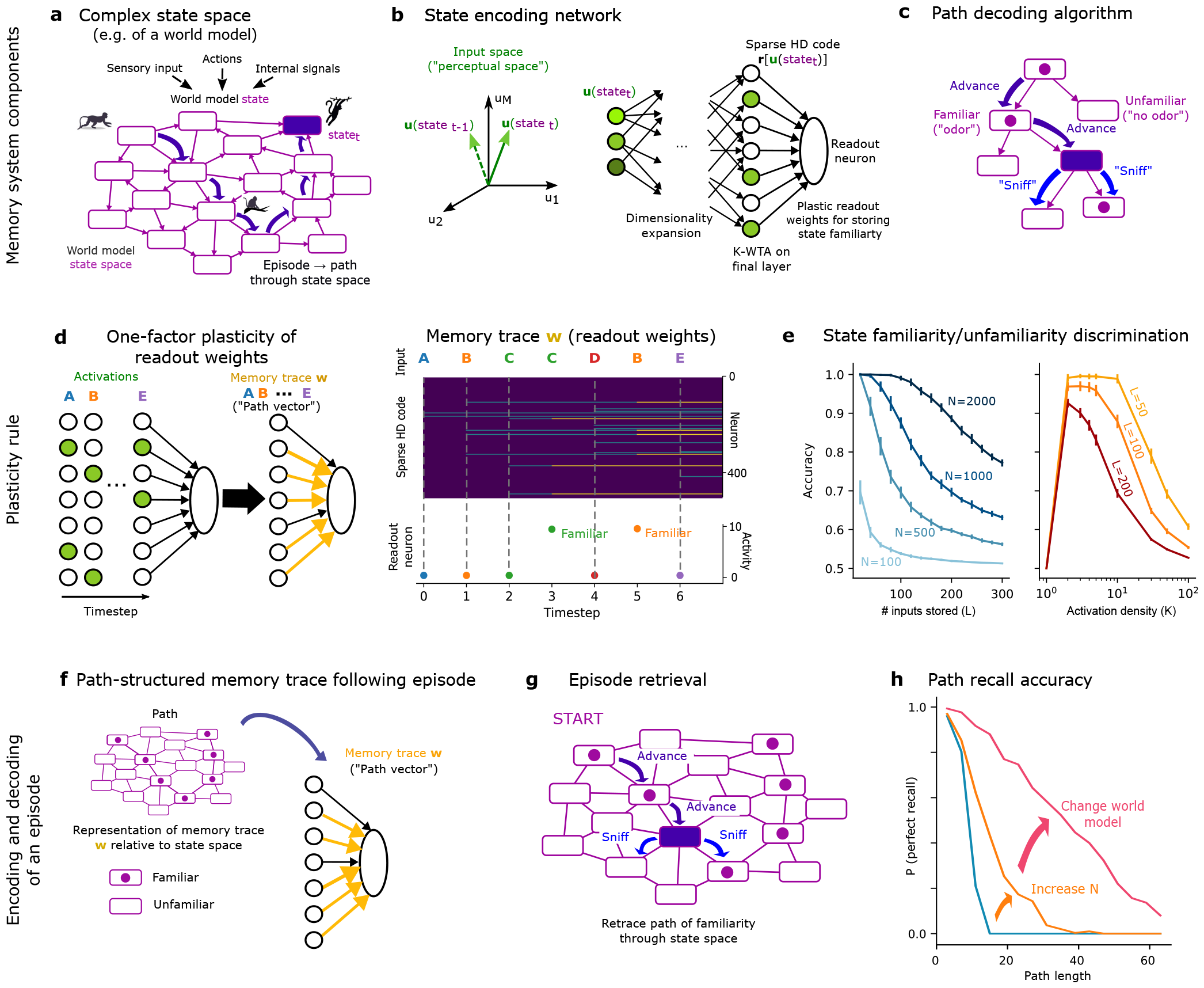
Episodic memory system using a one-factor plasticity rule. a) Schematic of a world model state space. b) Encoding network schematic. c) Schematic of odor-tracking-like decoding algorithm. Decoding occurs by following a path of familiarity (dots) through the state space. d) Left: Schematic of plasticity rule that creates the memory trace in the familiarity detector. Every time a neuron in the final layer activates, its synaptic weight onto the readout neuron increases by ^*η*^. Right: example evolution of the readout weight vector throughout the presentation of an input sequence and response of the readout neuron over the same input sequence. For this simulation ^*N* = 500, *M* = 50, *K* = 10, *N*_*layer*_ = 3, *G*_*in*_ = 10,*G* = 20,*q* = 0.1^, with each input represented as a random *M* -dimensional vector (Gaussian and i.i.d. across neurons). e) Familiar/unfamiliar state discrimination vs length of input (state) sequence ^*L*^ (assuming all inputs are unique), width of the encoding network (^*N*^), and activation density of the final layer (^*K*^). In left panel ^*K* = 30^, in right panel ^*N* = 500^. Here, ^*θ* = *K −* 1^; other parameters as in d. f) The final memory trace and its interpretation as a connected path through the world model state space. After a state sequence, the readout weights encode the set of states visited, which in the example shown trace out a unique path through the world model state space (dots represent familiarity). g) Schematic of retrieving an episode. h) Recall accuracy vs path length for three different simulations. Blue: ^*N* = 300^, orange: ^*N* = 2000^. Blue/orange: *N*^*world*^ = 1000,*K* = 10,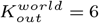 = 6. Green: ^*N* = 2000,*K* = 10,*N* *world*^ = 3000, 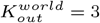 = 3, *N*^*world*^ is the number of world model states and 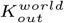 is the mean out-degree. The world model state space was constructed as Erdos-Renyi network with no self-connections, and ensuring at least one outgoing connection per state. Probability of perfect recall was estimated over 300 randomly sampled paths.

#### Plasticity rule

The only plastic component of our model is a single vector of weights ***w*** *∈* ℝ^*N*^ from *N* presynaptic neurons onto one readout neuron. (This will correspond to synapses from the final layer of the encoding network onto a readout neuron; see below.) Throughout our work, for simplicity, we will focus on a specific one-factor learning rule— long-term potentiation (LTP) triggered solely by presynaptic activity. As detailed later, however, other one-factor rules similarly capture our main results.

Let ***r***_*t*_ be the activity vector of the *N* presynaptic neurons at time *t*. The plasticity rule of the readout weights is simply:

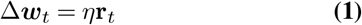

which states that the synaptic weight increase at time *t* is proportional to the current presynaptic activity via a learning rate *η*. Thus, at time *t*:

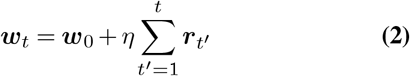

In other words, every timestep that neuron *i* activates, its synapse onto the readout neuron potentiates and remains potentiated for the course of the simulation; if it never activates during the simulation its synapse remains negligibly small (***w***_0_ = 0). For simplicity we do not include an explicit forgetting mechanism to prevent all synapses from potentiating without bound. Instead we reset ***w***_0_ to **0** in every simulation. The operating regime of our model is when *N* is substantially larger than the total number of potentiated synapses. In Fig. 3a - Fig. 3e we consider a continuous-time spike-based version of this rule, but everywhere else we use the discrete-time equations Eq. (1), Eq. (2).

**Fig. 3.**
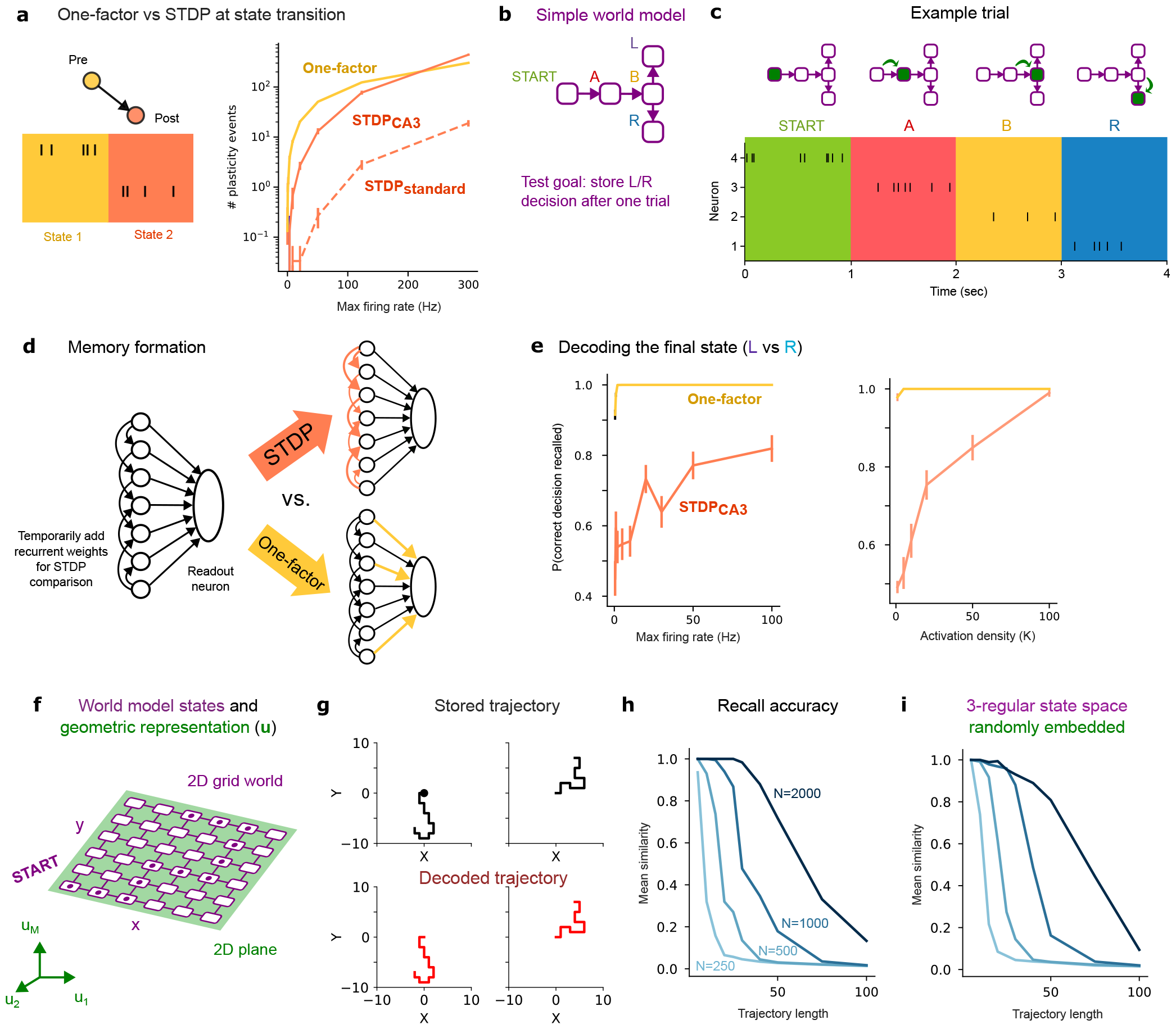
Comparison of one-factor plasticity vs STDP in the face of spiking, and recall of example world model trajectories. a) Left: Schematic of plasticity event counting simulation. Right: number of plasticity events surrounding a state transition for either the one-factor rule, a symmetric STDP rule based on CA3 (16), or a simplified standard STDP rule (10) including only potentiation, as a function of the max firing rate the pre- and postsynaptic cell can achieve. b) 5-state world model used for STDP vs one-factor comparison. c) Schematic of one trial. d) Schematic of memory storage via either STDP or a one-factor rule. e) Probability of recalling the correct decision after a single trial after storing the trajectory via either a one-factor rule or the CA3 STDP rule, as function of either max firing rate or activation density. f) Example world model and transitions supporting trajectories through a 2D grid world. Example path of familiarity is indicated with dots inscribed within states. g) Two example trajectories stored (independently) and recalled using a path vector. During the decoding/recall phase we introduce a heading variable (E, N, W, or S) to the agent exploration and prevent the agent from backstepping during recall. h) Similarity (1 minus edit distance) between stored and decoded 2D trajectory vs trajectory length and ^*N*^ . i) As in panel h, except for trajectories sampled from a 5000-state world model with a 3-regular directed graph structure (all states had 3 randomly picked downstream states).

#### State space

The next key ingredient is a complex state space that can be represented as a directed graph. A canonical example, which we will use throughout this work, is the state space underlying a *world model* (Fig. 2a)—an internal representation of an agent or animal’s environment commonly invoked for its utility in prediction, inference, and planning (32–36). In mammals, world models are thought to include complex state spaces that are acquired progressively (e.g. via gradual Hebbian plasticity) in neocortex (33, 37–41). Four key properties of such state spaces are relevant to our memory model: (1) states and their transition statistics can be represented as a directed graph—a standard approach in Reinforcement Learning (RL), for example; (2) states evolve in response to a variety of signals but can also be maintained without continued sensory input (e.g. via an attractor network (42–45)); (3) they enable simulation of state-action sequences; (4) the state space is not significantly plastic over the rapid timescale of episodic memory formation, i.e. episodes are not stored by changing the world model. Here we assume the agent has direct access to such a state space or world model, rather than modeling it mechanistically (e.g. as an attractor network), because our key results do not depend on its particular physical implementation.

To use such a state space for our memory model we invoke a crucial assumption. During a typical or natural behavioral episode the progression of the episode evolves the state along *an unbroken path through the state space that does not collide with itself* (Fig. 2a); thus, the *set* of states alone specifies an unambiguous path through the state space. For example, consider a world model whose states represent an animal’s position in a large physical environment, such that the state evolves to track the animal’s position. If the animal takes a path from its home through its environment, and does not return near any visited locations a second time, the set of states visited will specify an unambiguous path, akin to a trail of breadcrumbs, from the animal’s home to a specific location. Thus, typical navigational episodes often satisfy this criterion. A consequent prediction we will investigate is that deviation from this criterion (e.g. paths with loops or “branch points”) introduces recall errors.

As we will show, world models described by generic graphs, as well as state spaces that do not directly correspond to a world model—such as learned embedding spaces, or even the world itself—are also compatible with our framework. We focus predominantly on world models, however, due to their straightforward interpretability and direct relationship to natural behavior.

#### Encoding network

To transform world model states into a format suited for memory we use a simple random encoding network. First, we assume each state of the agent at time *t* is represented by a unique neural activity pattern **u**(state_*t*_) *∈ 𝒰 ≡* ℝ^*M*^ . This corresponds to a “perceptual” or geometric signature of the state, e.g. with similar states having similar **u** (39). We then pass **u** as input to an encoding network that converts it to a sparse, approximately binary high-dimensional (HD) neural code **r**[**u**(state_*t*_)] *∈* ℝ^*N*^, with *K* nonzero entries (Fig. 2b). Crucially, we desire the HD codes for different states to be near-orthogonal with high probability (46–50):

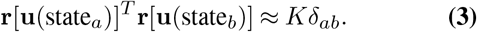

This follows from the action of the encoding network, which consists of random feed-forward layers and a final *K*-winner-take-all (K-WTA) operation (50, 51). Indeed, this network decorrelates or approximately orthogonalizes nearby patterns in *𝒰* (Fig. 2b, Fig. S1), recapitulating the traditional pseudorandom HD item representations used in human episodic memory models (49); sparse, decorrelated HD representations of odors in the insect mushroom body (52); and the recently discovered barcode-like representations of cache-locations in bird hippocampus (53).

To store visited states in memory, we apply the plasticity rule Eq. (1), Eq. (2) to synaptic weights ***w*** from the final layer of the encoding network onto a single readout neuron. At an episode start we reset ***w***_0_ = **0**. Then, after an episode consisting of *L* states, the final weight vector is

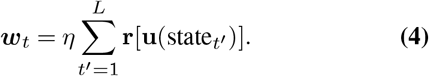

This trace ***w*** can also be naturally created online using Eq. (1); as the first several novel inputs activate largely non-overlapping sets of neurons in the final layer (provided *K ≪N*), the set of potentiated weights grows as new inputs are presented (Fig. 2d, right).

Functionally, ***w***_*t*_ enables familiarity detection of states, where familiarity means the state was visited during the episode. Let **r**_test_ be the HD code for a test state, and **r**_1_,…, **r**_*L*_ the HD codes for the *L* states visited in the episode. Let the readout activation be linear with weights ***w***_*t*_. Letting *η* =1 for simplicity, activation of the readout neuron, which we label 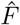, then indicates the familiarity of the test state:

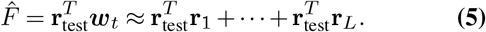

Using Eq. (3):

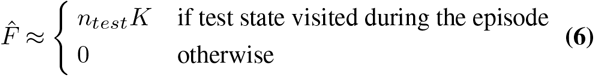

where *n*_*test*_ is the number of times the test state was visited. A threshold 0 *< θ ≤ K* can be chosen to binarize the familiarity detection. Accuracy of discriminating familiar states is high when *N ≫ L* and when the HD codes **r** are sufficiently but not overly sparse (Fig. 2e). Results are similar for other one-factor rules, including when weights are restricted to be binary (Fig. S2), hence do not strongly depend on implementation details.

Thus, by converting states to HD codes, then summing these via the one-factor rule, familiarity of states visited in the episode can be detected. This is mathematically similar to human recognition memory models and vector symbolic architectures in which item states are stored as sums of random HD vectors (46–49), as well as a Bloom filter (54)—a data structure for set membership queries—and a recent study on its possible neural implementation (55). The key difference is that here we use a multi-layer random network to compute HD codes on the fly (only two layers were used in (55)).

We emphasize that ***w***_*t*_ does not explicitly store the state *order*, nor does it directly enable pattern completion, as in a Hopfield network (56). Reconstruction of the episode instead depends on the fact that the visited states encoded in ***w***_*t*_ trace out a path through the state space. Thus, the *set* of states stored in ***w***_*t*_ encode the path. We therefore term the memory trace ***w***_*t*_ a high-dimensional *path vector* (Fig. 2f).

### Decoding algorithm

To decode an episode, the agent retraces the path through the state space, following the familiarity signal (Fig. 2c). At every step of decoding, we assume the agent is in one state. It then simulates potential actions to available downstream states and estimates each state’s familiarity 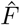 using the encoding network and ***w***_*t*_ (Eq. (5), Eq. (6)). If a state is familiar 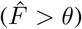 the agent advances to that state (if multiple downstream states are familiar, one is picked at random), then repeats this procedure until it reaches a state with no familiar downstream states. This resembles a simple odor-tracking algorithm, where an animal follows an odor trail by sniffing nearby locations and advancing when it detects the odor (57). Here the agent follows a path of familiarity through the world model state space. As with the world model, and following related work on repurposing chemotaxis for general problem solving (8), we do not model the decoding algorithm mechanistically (e.g. as a neural network), but rather assume that such an algorithm is accessible in the brain, since many animals have exquisite odor-tracking capabilities (57). In spatial tasks we also let the agent have a “heading” variable such that the previously visited state (behind the agent) is not considered to be downstream.

This algorithm is efficient. While retrieving the familiar states directly from ***w***_*t*_ would require testing all possible world model states, here due to the restricted transitions only states downstream of the current state need to be tested for familiarity. Thus, retrieval time is limited not by the size of the world model but by the out-degree of the familiar states.

### Complete storage and retrieval of an episode

The complete storage and retrieval of an episode occurs as follows. Before the episode begins, we put the agent in a specific “start” state, and we set ***w***_0_ to **0**. As an episode unfolds, the state evolves from the start state along a path through the world model that represents the episode. As each state is visited, its HD code **r** is created via the encoding network, which in turn updates the path vector ***w***_*t*_ via the one-factor rule (Eq. (1)). By the end of the episode the path is stored in ***w***_*t*_. To probe the ability of the agent to recall the episode we place it again in the start state, then let it sequentially reconstruct the episode by retracing the path via the decoding algorithm (Fig. 2g).

The essential result is that if 0 *< K, L≪N* and the visited states specify an unambiguous path, the path vector ***w***_*t*_ stores the path perfectly with high probability, and allows efficient decoding from the start state. Notably, the path can be a never-before-seen route through the state space—if the world model is large and complex the path can in turn be richly informative, while automatically exhibiting a natural sequential organization. Thus, a one-factor rule can store a complex sequentially structured episodic memory corresponding to a path through a state space like that of a world model.

Importantly, reconstruction is enabled but also constrained by the state space. While increasing *N* allows more states to be stored, the memorability and expressivity of paths depend heavily on the structure of the world model or state space (Fig. 2h). Intuitively, an ideal world model state space should admit many unambiguous paths that can each represent a unique behavioral episode, while also accurately reflecting the structure of the “world” that produces the episodes.

### One-factor plasticity is a robust alternative to STDP in the face of realistic spiking

Before examining the generality of path vectors we first demonstrate that for the purpose of storing familiarity, one-factor plasticity is a robust alternative to STDP in the face of realistic spiking. To this end we compare two alternative spike-based plasticity rules, operating at a synapse *w*^*\**^. The first rule, a spike-based one-factor rule is:

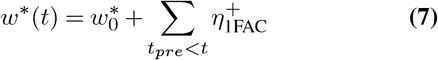

which increments *w*^*\**^ by 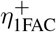 at every presynaptic spike time *t*_*pre*_, and 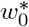 is the initial weight. The second rule, STDP, we define in terms of the difference 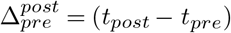 between pre- and postsynaptic spike times, operating via:

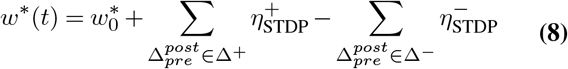

where the two sums run over all pairs of pre- and postsynaptic spike times prior to *t* that fall within Δ^+^ or Δ^*−*^, the potentiation and depression windows, respectively. Given a potentiating spike pair, *w*^*\**^ is increased by 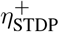 ; given a depressing spike pair, *w*^*\**^ is decreased by 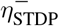. In general, both 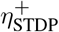 and 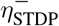 can be a function of (*t*_*post*_ *− t*_*pre*_) (58); however, for simplicity, we approximate such synaptic changes with rectangular functions. In both rules, one-factor and STDP, we bound *w*^*\**^ by 1. To be as conservative as possible in our comparison, we also initially let 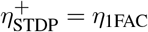 and 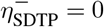, even though realistically we expect STDP to be substantially weaker (5, 15) and for the depression component to hinder potentiation.

One basic difference between our spike-based one-factor rule and STDP is the number of plasticity events (synaptic weights updates) mediating each mechanism. Whereas each one-factor event is just a presynaptic spike, each STDP event is a coincident pre- and postsynaptic spike pair. To highlight the importance of this difference, we considered a single synapse and the transition between two states, each visited for 1 second with no overlap. In the first state we let the presynaptic neuron emit Poisson spikes at rate *r*_max_, with the postsynaptic neuron silent; in the second state the presynaptic neuron was silent and the postsynaptic neuron emitted Poisson spikes at rate *r*_max_ (Fig. 3a, left). This mimics a simple scenario of a state transition during a task where the two neurons have different activity levels in the different states. Counting the number of plasticity events triggered by each rule, we found that the one-factor rule produced more plasticity events over a large range of firing rates, with STDP only exceeding the one-factor rule when the firing rate was greater than 100 Hz, and only for a specific form of STDP (based on CA3-CA3 synapses (16)) with a large symmetric coincidence detection window Δ^+^ = [*−* 100, 100]ms (Fig. 3a, right). Thus, when a pre- and postsynaptic neuron fire sequentially over a state transition, the one-factor rule will detect more plasticity events than STDP at realistic low firing rates, since it does not depend on coincident spiking.

We next compared the spike-based one-factor rule vs STDP in storing a simple episodic memory. Here we used a simple T-maze-like world model (Fig. 3b, top), with the goal of storing and decoding either of two possible trajectories (corresponding to a final left [L] or right [R] turn). During the episode storage, the agent spent 1 second in each state. Here we used the encoding network to transform each state into its sparse HD activity pattern **r**, then converted the resulting final-layer neural activations into Poisson spikes (Fig. 3c) at rates *r*_*max*_**r**, where *r*_*max*_ (Hz) was the maximum firing rate (since 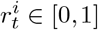. To compare one-factor vs STDP storage of the state sequence, we temporarily modified our encoding network. The modified network was identical to the original network (Fig. 2b) except that in addition to the readout weights ***w*** we also added sparse recurrent synaptic weights *W*_*rec*_ among the neurons in the last layer (Fig. 3d). We then compared two scenarios: in one case spiking activity induced STDP in *W*_*rec*_, while in the other it triggered the one-factor rule in the readout synapses ***w*** (Fig. 3d). We focused on the CA3-based STDP rule (16), which has a symmetric coincidence window of approximately 100 ms, since it produced more plasticity events than traditional STDP. This analysis can be seen as analogous to storing the memory either in the recurrent excitatory weights of CA3 (59) through STDP vs in the readout weights from CA3 to a single CA1 neuron through our one-factor rule.

For each condition we used a single trial (one-shot learning) to encode and store the trajectory. For simplicity we only sought to decode the final left/right decision *ŷ*. We decoded this either from the STDP trace *W*_*rec*_ via

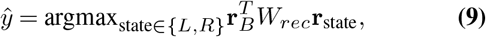

where **r**_*B*_ is the neural activity of the state before the decision, or from the one-factor trace (path vector) ***w*** via

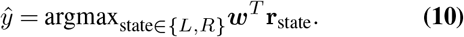

Intuitively, the STDP-based decoder uses the updated weights to make a prediction 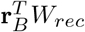 of what state followed B, then compares it via a dot product to either **r**_*L*_ or **r**_*R*_. The one-factor-based decoder reflects testing whether **r**_*L*_ or **r**_*R*_ was more familiar given ***w***.

In all of our simulations we found that the one-factor rule was as or more reliable than the STDP rule in storing the decision in our task. The one-factor trace retained the decision better than the STDP trace for all values of *r*_*max*_ we explored (Fig. 3e left); the STDP trace only achieved the same performance as the one-factor trace when the activation density *K* of the HD neural code was quite high (Fig. 3e, right). The one-factor trace also matched or outperformed the STDP trace upon varying *N* or the recurrent connection probability (Fig. S3).

Thus, the one-factor rule reliably stored a decodable memory of the final decision across a wider range of parameters than the STDP rule, suggesting its robustness in particular at low firing rates, sparse activation, and sparse connectivity, ultimately due to its indifference to spike-timing precision. While it remains an interesting open question whether STDP rules in the proper network context could significantly modify network function after a single episode, our results overall suggest that one-factor rules could be a robust, powerful, and potentially more biologically plausible alternative.

### Path vectors support diverse mnemonic tasks

Next, we show how the path vector representation can support a variety of one-shot recall tasks by leveraging the topologies of the world model and other state spaces. As we have established the robustness of the one-factor rule in the face of spiking (in addition to its likely faster and stronger operation than STDP), and to focus on the generic coding properties of the path vector rather than specific implementation details, in the rest of this work we return to using Eq. (1) and Eq. (2) directly to store the path vector, rather than spike trains.

#### Path vectors store trajectories through spatial and non-spatial world models

We tested our model’s ability to store and recall trajectories through a 2D open environment. The corresponding world model was represented as a plane embedded in the input space 𝒰 (Fig. 3f). To evaluate our model, we let the agent follow a random, non-self-intersecting trajectory of length *L*, which was then stored in the path vector ***w***_*t*_. We evaluated recall by placing the agent back at the origin and applying the decoding algorithm. The decoding algorithm only tested candidate next states to the left, right, and ahead (but not behind) of the agent’s current position. In general, such trajectories were successfully stored and recalled by the model (Fig. 3g), with recall accuracy decreasing with the length of the trajectory and increasing with *N* (Fig. 3h). The path vector can thus store episodes corresponding to non-intersecting paths through 2D open environments, a fundamental type of memory required for one-shot memory of navigational routes through physical spaces.

We also found that path vector code reliably stores trajectories through non-spatial world models. We created a world model with 5000 states and a 3-regular directed graph structure (each state has three downstream states). Each state was represented by a random 50-dimensional input vector **u**_*state*_ reflecting a unique “perceptual signature” associated with the state. We sampled random trajectories starting from a chosen start state, stored them in the path vector, and attempted to recall them from the start state. As with the spatial world model, trajectories through the 3-regular world model could be recalled via the path vector. Thus, our mechanism can operate on generic world models, as long as *N* is large enough to adequately discriminate familiarity and the path can be reconstructed from the set of visited states. Indeed, recall is impaired when the world model does not match the transitions used to produce the input state sequence (Fig. S3).

#### Serial order and arbitrary associations can be stored as paths through learned embedding spaces

We have demonstrated encoding and decoding of sequences whose adjacent elements correspond to adjacent world model states. This reflects our assumption that typical or natural episodes trace out paths through a world model (Fig. 2a). But what if the sequence doesn’t follow such structure? Can a path vector encode arbitrary sequences of inputs like a sequence of items presented in a serial recall task (49)? In our model, this problem is solved by mapping such sequence to an unambiguous path, either through part of the world model or through another state space like a learned embedding space (60), of dimension *m* and indexed by coordinates ***z*** = (*z*_1_,…, *z*_*m*_) (in our simulations we use a discretized space). The key property of such a space is that certain states explicitly represent items but many or most states do not. Instead, these “auxiliary” states can be used to construct paths through the embedding space that connect the item states in order (Fig. 4a). Recall is achieved by retracing the path and reporting item states visited along the way. In Fig. 4b we show an example of recall/decoding in this setup in which the path is unambiguous. By following the path through the embedding space the four items A-C-B-D are revisited in the correct order.

**Fig. 4.**
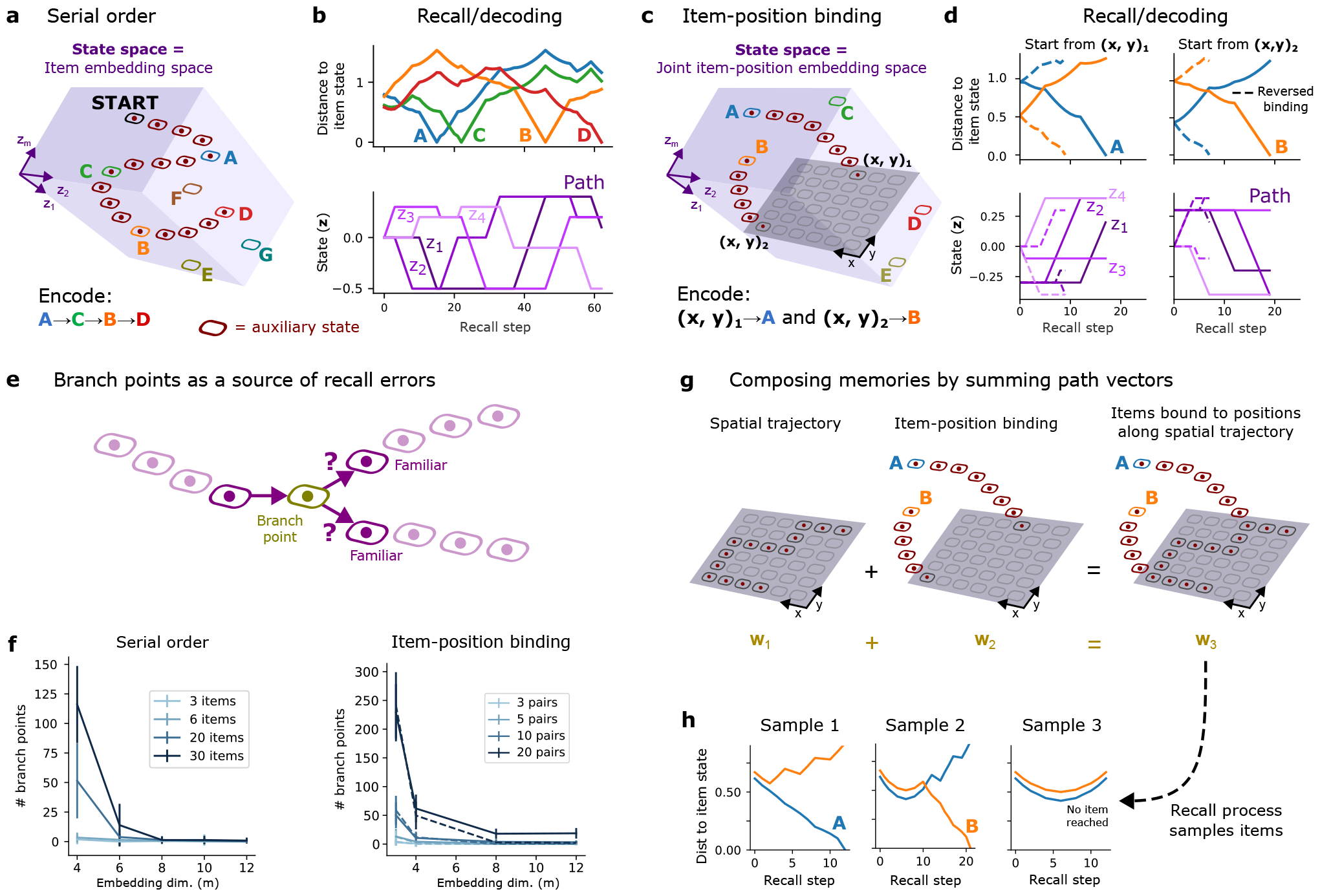
Path vectors applied to embedding spaces support diverse memory tasks. a) Path-vector scheme for storing serial order. The rest of the auxiliary states in the embedding space, which are not included in the path, are present but for visualization purposes are not shown here. b) Example decoding of serial order by following the path through the auxiliary states in the embedding space. c) Path-vector scheme for binding items to spatial positions via a joint item-position embedding space. d) Summary of decoding which items are bound to which locations by starting at the position states and following the path to either item. e) Schematic of emergence of branch points. f) Number of branch points vs embedding dimensionality for serial order (left) or item-position binding (right) example. g) Schematic of composing memories by summing path vectors created in overlapping state spaces. h) Examples of different outcomes of applying the odor-tracking algorithm to the composite memory, starting at the same position on the grid.

Note that we do *not* predict path vectors to be the primary memory substrate used in immediate serial recall tasks. For instance, sequential information could also be temporarily stored in the activity pattern of a recurrent neural network (43, 61). Our claim is instead that the path vector mechanism *could* be recruited for such a task (intuitively, using episodic memory to solve a working memory task), likely also extending the lifetime of the memory by engaging LTP. In fact, mapping an arbitrary sequence to a unique path through a world model or other state space may have a stronger tie to the well-known mnemonic trick of building an auxiliary story to remember longer lists of words (62), or linking items through evocative mental imagery (63), if one associates the story/imagery with creating paths through a specific state space, making the list easier to recall. Thus, our one-factor rule can be used to form an episodic memory encoding arbitrary serial order, conditional on the ability to construct a path satisfying the desired order among the items.

A second example use case of auxiliary states within an embedding space is to store pairwise associative memories. For instance, consider the task of remembering which items are present at different locations of a 2D space (64). In our model this task can be solved by considering a multidimensional item embedding space, in which a 2D plane representing positions is itself embedded (Fig. 4c). (As our model is compatible with a vast set of states, the apparent “curse of dimensionality” of such product spaces, which could have been learned over a lifetime, poses no issues, since the path vector still only stores a small subset of these.) To bind items to positions, a path is constructed through this space from each position state to its associated item state (Fig. 4c). Starting at a position state and using the decoding algorithm to follow the path of familiar states through the embedding space then leads to the corresponding item state (Fig. 4d), as long as the paths linking each location to each item do not collide.

Thus, path vectors may serve as a general-purpose one-shot, one-factor memory representation that can support not just world model paths, but generic serial and associative structures, provided that an appropriate state space is available and valid paths can be constructed.

#### Branch points are a path-specific source of recall errors

Our model admits a specific type of errors when representing information as paths. We term a state in the path with more than one downstream familiar states a “branch point”. This can occur if the path intersects or “collides” with itself (comes within one network link of the existing path), introducing ambiguity and potential errors in recall (Fig. 4e). At the branch point, there is no unique state to visit next, so the decoding algorithm can revisit states out-of-order. The probability of branch points occurring depends on the structure of the state space and how the path is built. In the serial recall and position-item binding tasks, paths linking items or position-item pairs are much more likely to collide when (1) the dimensionality of the embedding space is low, and (2) more items are stored (Fig. 4f), inducing more branch points and recall errors. Thus, branch points can be limited by employing paths through higher dimensional embedding spaces.

#### Summing path vectors stores and composes multiple memories

Path vectors operate by aggregating a collection of states into a common representation encoding an episodic memory, but this can be generalized to the aggregation of multiple episodic memories together. Whether memories interact or not depends on the states they share. When two path vectors ***w***_1_ and ***w***_2_ are summed into a composite path vector, the result ***w***_3_ = ***w***_1_ + ***w***_2_ encodes the union of the states encoded in each path (when *K, L*_1_ + *L*_2_ *≪ N*). We note that this sum is also enacted naturally by the same one-factor rule Eq. (2) used to aggregate states into a single path vector. In general, paths through distinct state spaces won’t interfere. Therefore, recalling from either of the path vectors is equivalent to recalling from the composite path vector. The same *N* neurons are thus used to store multiple episodic memories supporting different tasks, instead of each task requiring a separate memory “slot”. This accords naturally with the use of a common memory substrate to encode different tasks that might unfold in different contexts, spaces, or sensory modalities, a role often ascribed to hippocampus (59).

When the states stored in the two path vectors overlap, the composite (summed) path vector can produce interference that yields novel recall patterns. For example, the sum of a path vector ***w***_1_ encoding a path through 2D space with a path vector ***w***_2_ encoding two items paired with two positions along the 2D path (Fig. 4g) encodes the union of the states stored in ***w***_1_ and ***w***_2_, which defines a branching path through the full state space (Fig. 4g). Recalling from this path vector yields trajectories that stochastically visit each item state (Fig. 4h), effectively sampling items using the composite path vector. Thus, under the path vector formulation, episodic memories can be combined to produce compositional path vectors that may be useful for computations beyond recall alone. The path vector can be used, for instance, to hold in memory and sample multiple alternatives, a fundamental computational primitive for inference.

### Policies can be learned by summing episodes’ path vectors together contingent on reward

Finally, we asked how path vectors could be used for reward-driven learning, which is essential for the brain’s ability to learn tasks and adapt to novel environments. We expanded the 2D navigation model discussed earlier to function in a reward-based environment. Imagine an agent, or animal, navigating a grid world to earn a reward. We let the starting location of the agent vary across trials, while fixing the reward location (Fig. 5a). In trial *n*, the agent takes a path through the environment, which it stores in a temporary path vector ***w***_*n*_. Over multiple trials, the agent converts these into a second representation **p** *∈*ℝ^*N*^, created via a nonlinear weighted sum of the path vectors corresponding to paths that led to reward (Fig. 5b). One can think of each ***w***_*n*_ as a temporary episodic memory associated with trial *n*, and **p** as a permanent reward-dependent representation of the environment formed by summing (element-wise) the rewarded path vectors together. Biologically these could be implemented by two different one-factor rules operating on fast and slow timescales, but in the same synapses. Intuitively, the environment representation ***p*** encodes a map of effective familiarity levels of the states, which can be retrieved via the dot product ***p***^*T*^ ***r***(state_*i*_).

**Fig. 5.**
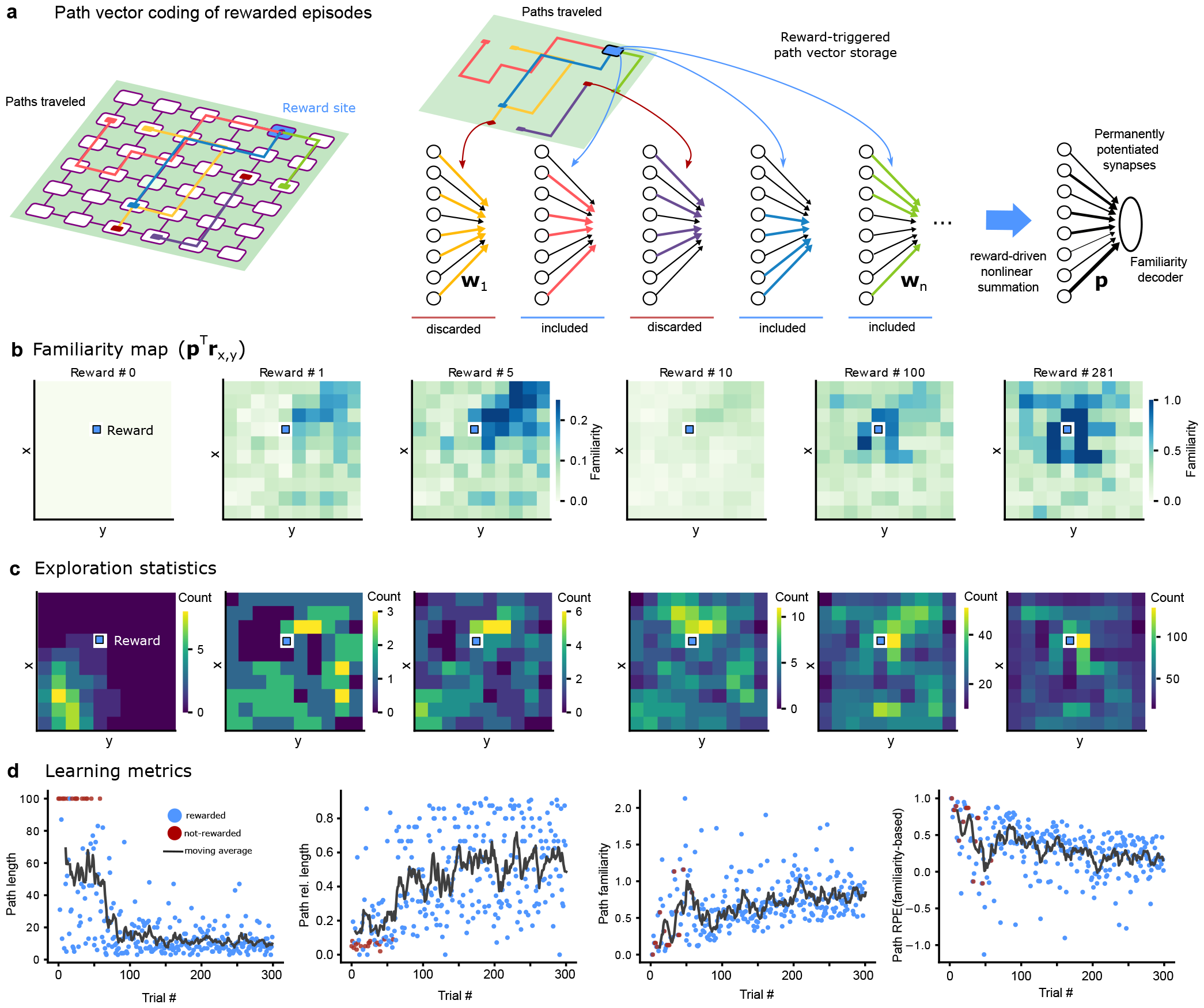
Learning with path vectors. a) Left: Schematic of paths traveled in rewarded and unrewarded episodes. Right: cartoon representing included and discarded paths in the reward-driven nonlinear summation generating the path vector familiarity decoder. b) Familiarity maps during learning. For each location in the environment effective familiarity can be decoded through the learned path vector familiarity decoder. As learning proceeds, locations closer to the reward become increasingly more familiar. c) Statistics of explored locations during learning. This is one of the leading factors that drives learning. d) Learning metrics. Left: Path length corresponding to the total amount of steps taken by the agent in the environment during the current trial. If the path traveled reaches 100 steps the trial ends without a reward. The black line displays a moving average over the previous 10 trials. Center-left: Path relative length is the shortest path length leading to the reward from the initial location of the agent, divided by the length of the path taken by the agent. Center-right: decoded familiarity for the current trial path. Right: Familiarity-based path Reward Prediction Error (RPE). Learning follows a scheme akin to Reinforcement-Learning algorithms driven by an RPE signal, except here we use familiarity-based RPE. While this panel shows the same data as the previous one, it emphasizes the decreasing RPE over learning.

In our simulation we let an agent navigate a 10×10 grid world. Prior to the simulation **p** was initialized to be near zero. The trajectory at each trial was generated via the policy:

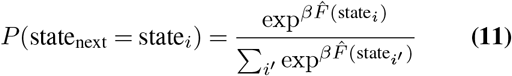

with *i* indexing all downstream states from the current state, and 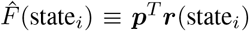 indicating the familiarity of state *i* with respect to ***p***. The parameter *β* controls the noise in the process. For large *β* the agent always selects the most familiar state, whereas for intermediate *β* state probabilities follow their relative familiarity. Trajectories were truncated either when reward was received or after 100 timesteps.

Over multiple trials, we let the agent learn a policy by updating ***p***. On trial *n* all weights ***w*** are set to 0. The temporary path vector ***w***_*n*_ is created after the agent traverses a route through the environment. Next, the “path familiarity” 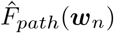 is computed via:

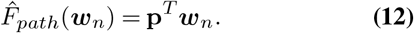

If no reward is received ***w***_*n*_ is discarded, and the agent moves onto the next trial. If reward is received, however, **p** is updated according to

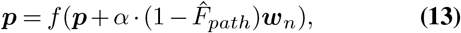

which in turn updates the policy Eq. (11). Similar to traditional RL models, 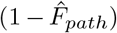 plays a role akin to reward-prediction error and *α* is the learning rate. We used *f* (*x*)= tanh(*x*) applied component-wise to bound components of **p** at 1. This is similar to the previous section in which memories were composed by summing path vectors (Fig. 4g) except here summation is conditional, weighted, and nonlinear. Formally, this is a reward-mediated (i.e. two-factor) learning rule, except that rewards simply function as a mechanism to trigger permanent plasticity and carry no information about either the identity of the synapses to modify or how they should change—it is commonly believed that dopamine or other neuromodulators could provide such a signal (15). The crucial difference between our learning rule and traditional RL algorithms (32) is that here the entire trajectory, when rewarded, is stored, simply by (weighted) adding of the latest path vector to ***p***. This framework is similar to Q-learning reinforcement learning algorithms. However, instead of using a value function, it uses a familiarity (65) map built out of paths.

Over the course of our simulation, the agent progressively learned a representation ***p*** of the environment (Fig. 5b). After a handful of trials locations (states) closer to the reward started acquiring a higher familiarity. The paths leading to the reward location often intersect, resulting in overlapping state statistics in the corresponding path vectors. After being rewarded a few times, the agent learned to reach the reward in just a few steps (Fig. 5d). We note that the number of steps is not fully optimal as the agent tends to reach the reward by repeatedly following similar trajectories. We found these trajectories to be quasi-optimal, being only 20-30% longer than the shortest possible path. As expected, the reward prediction continued to improve throughout the learning process. Finally, we note that these highlighted properties are crucially dependent on the agent’s exploration being tied to the policy throughout the learning process (on-policy scenario). The off-policy scenario where exploration is separated from the computation and exploitation of the familiarity map is illustrated in Fig. S4.

Altogether, these results show that path vectors are a feasible and efficient, and by our previous arguments more biologically plausible, basis for reward-mediated policy learning.

## Discussion

Our study emphasizes the potential significance of one-factor plasticity in memory and learning systems. This rule represents a robust, biologically plausible alternative to STDP for one-shot learning that can create highly expressive memory traces. Ultimately, the capabilities of our proposed system derive from summing high-dimensional (HD) vectors (46, 48, 54, 55) to store subsets of states sampled from a pre-existing structured state space. This in turn allows the rule to store complex episodic information as expressive, information-rich paths through that space that can be efficiently decoded. This model thus serves as a comprehensive alternative to Hebbian theory, aligning with the broad observation that biological plasticity rules with timescales that match one-shot memory formation are largely non-Hebbian (21, 24, 26–29). Our work thus poises one-factor and related non-Hebbian rules as significant tools in the field of memory systems. Our work provides biophysically grounded theoretical support for the representations of episodes (30, 36, 66) and policies in memory as paths (or collections of paths) through a rich, pre-existing state space. This has at least three broad implications. First, our model reveals a natural interpolation between episodes and policies. In combination with the decoding algorithm, even a single episode immediately specifies a policy, guiding the journey to a reward site from various starting states (even generalizing to unfamiliar start states, which the policy exits via random search). However, more efficient policies can be made from multiple rewarding episodes simply by summing their path vectors together (Fig. 5). In contrast, most existing models treat one-shot episodic memory as a generic buffer for temporarily storing arbitrary information (6, 7, 34, 36, 39, 41, 49, 67, 68) (typically through Hebbian plasticity), with policies existing in a separate function space. In episodic RL, for instance, episodes are stored as memorized action sequences that can be sampled to efficiently approximate state-action values to shape policy (34, 36). Our work similarly champions the utility of episodes for efficient policy learning, but differs from the prevailing episodic RL framework by skipping the construction of a value function and instead forming policies by directly summing episodes together, which still leads to efficient reward finding (Fig. 5d). Our model thus proposes a slightly simpler, more biologically plausible version of episodic RL. Moreover, states, episodes, and policies under our model are consequently all represented in the same vector space, ℝ^*N*^, revealing a potentially useful format for other, more complex computations on these objects.

Second, our model posits that the structure of the world model should impact recall. Because our proposed memory system operates relative to the state space of the world model, episodes encoded as unambiguous paths are easier to retrieve. This suggests that individuals with different world models may remember the same episode differently, in line with reconstructive and schema theories of memory derived from human experiments (69, 70), which are highly relevant to memory reconstruction in critical real-life settings like witness testimony (71). Our work is fully consistent with these theories, suggesting that using schemas for reconstruction is akin to traversing a path through a complex state space, which could have rich structure learned over a lifetime (there is essentially no limit to the size or complexity of the state spaces compatible with our model). Our work thus extends existing theories of reconstructive memory by establishing consistency with more biologically realistic plasticity rules, and by emphasizing the role of the state space topology. Our model in turn suggests high-level explanations of complex phenomena, such as coherent narratives being easier to recall than scrambled ones (72)—we predict the former can be more readily encoded as clear paths through a world model. This in turn hints at a mechanism underlying the mnemonic device of enhancing serial recall by imposing auxiliary narrative structure (62), which we suggest reflects encoding an item list as an unambiguous path.

Third, path vectors particularly benefit one-shot learning in large, novel state spaces. Because the HD codes **r** are computed on the fly (Fig. 2b), state representations and connectivity need not be learned. A path vector can thus represent an episode as a path through a completely novel environment (whose state transitions are constrained by physical adjacency), which in combination with an odor-tracking-like algorithm acts like a one-shot policy. Our work thus supports and extends the recent “endotaxis” model, which showed how odor-tracking could be repurposed for one-shot learning of goal-oriented tasks (8). In endotaxis, an agent learns state connectivity via Hebbian plasticity, along with artificial gradients to goal sites, which it then ascends using a chemotaxis-like algorithm (57). In contrast, our model does not require rapid Hebbian plasticity to learn state transitions, suggesting a simpler, more bioplausible solution. Moreover, following a path vector only requires estimating a binary familiarity signal, which could be faster to estimate from stochastic spikes than a gradient. Functionally, our model bears a stronger resemblance to theories of landmark navigation in insects, where it was proposed that odor tracking algorithms may be repurposed to follow generic attractive signals like familiarity (73). Thus, our work sheds new light on simple, biological ways in which ancient olfactory or foraging algorithms (74) could be repurposed for generic tasks.

### Predictions

Our model generates a number of testable predictions. First, impairing Hebbian plasticity should not degrade one-shot episodic memory. This could ideally be tested in naturalistic tasks such as single-trial route learning (75), where we predict that acute pharmacological blocking of STDP would not impair learning, so long as other plasticity mechanisms were not degraded.

Second, episode recall should follow the pre-existing state space transition structure more reliably than the temporal structure of the original experience. This is supported by observations in rodents where hippocampal replay of recent routes followed a roughly uniform speed (8 virtual m/s) independent of the temporal structure of the original trajectory (76). A more specific version of the prediction could be tested in virtual reality settings (77) by training mice to learn a virtual maze, then presenting a sequence of maze locations out-of-order. We predict that replay of these locations will align with their topology in the maze rather than the sequence of their presentation. A similar prediction could be tested in humans by teaching subjects transition statistics within a complex artificial state space, then probing recall for sequences in and out of alignment with the state space.

Our final prediction is that the recall of complex input sequences—such as a movie, which in our model would be encoded as a complex path through a world model—will be impaired if the same world model state is revisited during the encoding. This procedure would generate branch points (Fig. 4e) that degrade the retrieval performance. This could potentially be tested via neuroimaging, if world model states have reliably measurable neural representations. We anticipate an increase in recall errors surrounding world model states visited multiple times during the encoding.

### Limitations and synergies

Although our model has significant implications, it is also subject to certain limitations that motivate further investigation. We briefly examine three areas, highlighting the limitations of our work and its synergies with more nuanced biological non-Hebbian rules.

First, our model can only store familiarity of *𝒪*(*N*) states (see Supplement). Thus, the number of episodes that can be reliably stored depends on the number of states visited per episode and the degree of overlap/adjacency of their corresponding paths. To illustrate, consider a scenario where the one-factor rule can store *cN* distinguishable states (*c<* 1). In this case, the maximum number of episodes of length *L* that can be stored is approximately *cN/L*. If the set of episodes subsequently specifies *cN/L* unambiguous paths, each episode can be fully reconstructed from its initial state. Note that our setup differs from traditional memory models (e.g. attractor neural networks) that store individual inputs (43, 56)—in our model episodes contain sequences of states, and recall depends on the graph structure of the state space and the alignment of the paths with that structure in a complex way. However, similar to existing theories, we also expect that important episodes could be consolidated over time into updated world model statistics, thus freeing space in the episodic system (7, 39). Future work is warranted to directly compare our model’s capacity with existing benchmarks on a variety of specific world model structures, and in the context of additional consolidation processes.

The capacity limitation follows from our model’s reliance on a single readout neuron. However, naively incorporating multiple readouts may be counterproductive for two reasons. First, changing synaptic weights can be resource intensive. Therefore, if only a few states need to be stored (a task easily accomplished with one readout neuron), altering all readouts may be unnecessarily costly. Second, if each readout can dependably store *cN* states in *N* synapses, then when the total number of states is greater than *cN*, all readout neurons will become saturated, leading to catastrophic forgetting. An alternative strategy would be to allow plasticity to occur in only a select few readouts at a time. In fact, this is reminiscent of how BTSP of synapses onto CA1 cells from CA3 seems to function, with approximately 5-second plasticity windows in certain CA1 neurons opened by occasional dendritic plateau potentials (24), which in the context of our model would cause states to be stored in a different sparse set of readout neurons every several seconds. State familiarity could then be assessed by summing over all readout neurons: more readout neurons will activate for familiar states than unfamiliar states. (We note that if implemented in CA1 this would not rule out the possibility of novelty detection in other neurons (78).) Therefore, using different readout neurons at different times allows for the recognition of more states. This suggests a novel functional role of BTSP—an established yet puzzling plasticity rule—in enhancing the capacity of the already flexible memory system we have described. Exploring in more detail how our model synergizes with BTSP will be an exciting future direction.

A third opportunity to enhance our model is by including additional roles for dopaminergic neuromodulation. In the fly mushroom body, coarse dopaminergic inputs onto Kenyon-to-Output-neuron synapses shape when and how plasticity occurs (26, 27). While we have proposed dopamine as an error signal facilitating the transformation of path vectors into a permanent memory trace (Fig. 5), dopamine could also play a secondary role in the online gating of state storage (79) as the episode progresses. For instance, if the episode traverses a section of the world model consisting of a chain of states, there is no need to store the intermediate states of the chain, as they can be unambiguously recovered from the world model even without a familiarity signal, so fewer states need to be stored overall to represent an episode. Dopamine could serve as a natural mechanism for signaling the saliency of which states to store, since the most important ones to store are those that do not deterministically follow the upstream state. This approach would compress the path vector code, enabling it to encode longer paths in the readout weights.

### Open questions

Our model paves the way for a new set of inquiries. While our exploration has been limited to relatively simple world models and state spaces, it’s likely that the world models in higher mammals are considerably more complex. Two overarching questions that remain to be addressed include: how the structure of these state spaces and the strategies for converting episodes into paths influence recall, and how these structures and strategies can facilitate generic computational and learning problems. Moreover, it would be intriguing to establish a connection between the path vector model—which we believe is most relevant for storing natural episodes that are seldom studied in the laboratory—and neural concepts used to explain controlled memory tests, such as temporal context (80) and persistent activity in recurrent neural networks (43, 61).

## Materials and methods

### Encoding network

Encoding networks were multi-layer feed-forward networks. All layers had the same number of neurons *N* . The first layer mapped the *M* -dimensional input vector **u** to an *N* -dimensional activation **c**_0_ via

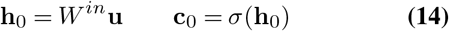

where *σ*(*h*)= 1*/*(1 +exp (*− h*)). All middle layer activations (with weights *W*^*d*^, where *d* indexes the layer) were given by

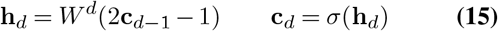

and the final (*D*-th) layer activation by

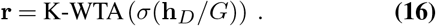

The input weight matrix had i.i.d. Gaussian weights with 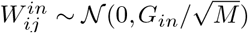 All middle layer weight matrices were sparse (with connection probability *Q*), with nonzero connections all i.i.d. and sampled as 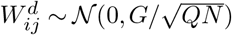 where *G*_*in*_ and *G* are gain parameters.

### Learning rate

When applying the plasticity rule Eq. (1), Eq. (2), we used a learning rate of *η* =1 unless otherwise noted.

### Input representations

For state spaces without spatial structure (e.g. Fig. 3i), input representations **u** *∈* ℝ^*M*^ of each state were given by *M* -D vectors with i.i.d. Gaussian elements. For state spaces with geometric structure (e.g. Fig. 3f) inputs were represented as constrained to a low-dimensional manifold (e.g. a 2D plane) embedded in the *M* -D ambient space. In such cases the embedding was created by picking an “origin” 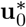, with 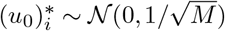 and *m* pseudo-orthogonal “basis vectors” 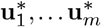 (where e.g. *m* =2 for a 2-D plane), with 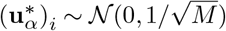 State coordinates **z** *∈* ℝ ^*m*^ (e.g. Fig. 4a) were then represented as

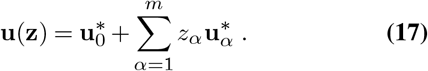

This formulation essentially builds an *m*-D hyperplane (with basis vectors random but near-orthogonal since *M* = 50) embedded in the *M* -D ambient space. The reason we did not simply use an *m*-D input space ℝ^*m*^ was because in that case coordinates **z** near the origin have low norm compared to co-ordinates far from the origin, which confounds the HD encoding **r** (the pattern separation functionality of the encoding network works best when inputs have similar norm). When an *m*-dimensional hyperplane is embedded in a space where *M m*, this is less of a problem, since different states’ input representations **u** all have similar norm. To further retain approximately uniform norms over the geometric state spaces, we constrained **z** *∈* [*−*1, 1] ^*m*^

### Spike-based simulations

Spike trains used in Fig. 3 simulations were sampled from a Poisson process with a state-dependent firing rate. In Fig. 3a, no encoding network was used; we simply simulated spikes in two synaptically connected neurons, by setting the presynaptic rate to *r*_*max*_ and the postsynaptic rate to 0 for 1*s< t<* 0 (state 1), and the postsynaptic rate to *r*_*max*_ and the presynaptic rate to 0 for 0 *<t<* 1*s* (state 2).

We used a simplified version of STDP capturing the main coincidence dependence. In our “standard STDP”, a plasticity event was counted as every occurrence of a postsynaptic spike following a presynaptic spike by less than 20 ms (we ignored the negative lobe of (58)). In CA3-inspired STDP, which has a symmetric window (16), a plasticity event was counted whenever a pre- and post-synaptic spike occurred within 100 ms of each other, regardless of order. Since the CA3 STDP rule gave more conservative results we used only this rule in our comparisons in Fig. 3e.

In Fig. 3e firing rates were determined using the encoding network. Each state was represented by a vector **u**, combining Eq. (17) with the geometry of the T-maze, using a step-size of 0.5 in **z**-space (**z**_start_ = (0, 0), **z**_*A*_ = (0.5, 0), etc.). HD codes **r** were created from state representations **u**, which were in turn converted to firing rates (in Hz) by multiplication with a scalar *r*_*max*_ (since the HD codes **r** were between 0 and 1 due to the sigmoidal nonlinearity). Spikes in each neuron were sampled from a Poisson process conditioned on these rates. During the simulation each state (START, A, B, and L or R) was visited for 1 second, and spikes elicited in each neuron according to its state-dependent activation.

For a given parameter set a simulation proceeded as follows. First an initial set of readout weights ***w***_0_ and recurrent weights 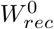 were created. The recurrent weights were sparse, with connection probability 0.02. All elements of ***w***_0_ and all nonzero elements of 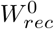 were then sampled i.i.d. from a gamma distribution with mean 0.05 and standard deviation 0.05, i.e. all nonzero initial weights came from the same distribution. Each simulation consisted of 50 trials. In each trial the readout and recurrent weights were reset to 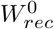 and ***w***_0_, then the task performed, spikes generated, and either the one-factor or STDP rule applied. Finally, the decision (L or R) was decoded using Eq. (9) (STDP) or Eq. (10) (one-factor). 100 simulations were used for each parameter set and means and standard errors for decoding accuracy reported in Fig. 3e, Fig. S3.

### Serial order and item-location binding paths

In the serial order example (Fig. 4b), 4 item states (A, B, C, D) were assigned random positions in a 4-D embedding space. In the item-location binding example (Fig. 4d), the first 2 dimensions corresponded to the plane representing 2-D physical space. Subpaths linking consecutive items (serial order) or item-location pairs were randomly constructed as follows. The embedding space (**z**-space) was an *m*-D lattice with spacing of 0.1. To create a subpath between two locations, first a random permutation of embedding space dimensions was created (excluding the first two dimensions corresponding to the plane in the item-location binding simulation). The subpath was then constructed by traveling along one dimension at a time (following the permutation order) until the target state (e.g. the next item) was reached. We used this algorithm instead of approximating diagonal shortest paths on a lattice in order to avoid spurious path collisions at kinks in the path caused by discretization.

### Alternative one-factor rules

Alternative one-factor rules for familiarity detection were also investigated (Fig. S2). These all yielded highly similar results for state familiarity detection because they operate under the same mathematical principle: storing a collection of *N* -dimensional vectors in a single *N* -dimensional memory trace purely via element-wise operations. We note that although synaptic vs excitability changes are often contrasted (28, 29), here one-factor rules, whether synaptic or excitability-based, essentially represent alternative implementations of the same computation.

The potentiation-based presynaptic rule was equivalent to Eq. (2), with *η* = 1, except that weights could not exceed a value of 1. In other words, a presynaptic neuron activating triggered its synapse onto the readout neuron to switch from 0 to 1 and remain potentiated throughout the simulation. The depression-based presynaptic rule (Fig. S2 c) was identical to the potentiation-based presynaptic rule (Fig. 2e), except that: (1) readout weights were initialized to 1, (2) readout weights from active neurons were set to 0 upon activation, and (3) familiarity was asserted if **r**_*test*_***w*** was *below* a threshold.

In the excitability-based rule (Fig. S2 d), all readout weights were fixed at 1. The activation level of neuron *i* in the final layer, however, depended on an excitability level *b*, which was increased whenever that neuron activated in response to a state input **u**. Intuitively, this causes neurons to exhibit an increased response to familiar inputs because their excitability increases during the original presentation of the input (28, 30). Specifically, the final layer activation (in contrast to Eq. (16)) was given by **r** = K-WTA(**h**_*D*_ + **b**). Note that this caused HD codes **r** to deviate from the binary approximation (because neurons had to activate to trigger the excitability change (28), but still had to be able to increase their response to a fixed input, since the readout weights remain fixed). Initially, we set **b** = 0, then set *b*_*i*_*→* 1 if *r*_*i*_ *>* 0 during the presentation of a state. The final familiarity readout was then 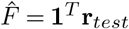, with values above threshold indicating familiarity and those below threshold indicating novelty.

The adaptation-based model (Fig. S2e) was like the excitability-based model except that *b*_*i*_*→ −*2 if *r*_*i*_ *>* 0, thereby decreasing the responses of neurons to repeated inputs. Familiarity was then determined if the value **1**^*T*^ **r**_*test*_ was below threshold.

### Learning rewarded episodes with path vectors

We generalized the one-factor learning scheme to operate in the presence of rewards, in the case of an agent navigating a 10×10 grid. The agent learned a path familiarity as described in the main text. Simulations for Figs. 5b to 5d entailed 300 trials with a fixed reward location, but a random initial location. The agent’s policy, captured by Eq. (11) had a single temperature parameter *β* that followed a simple annealing decay determined by *β* = *β*_0_N_*rewards*_*/T* with *β*_0_ = 10, *T* = 100 and fixed at N_*rewards*_ being the total number of accumulated rewards up to the current trial. The learning dynamics, captured by Eq. (13), used a single learning rate parameter *α* = .5. Results displayed in Figs. S4a to S4b follow an identical setup except that the agent’s behavior is off-policy—at every step the agent selects an action randomly (excluding the action leading to the last explored state). Therefore the parameter *β*, and more in general Eq. (11), are not relevant for this setup. Other values for this simulation are reported in Table S1. Reasonable changes in such parameters (e.g. *M* varying from 10 to 100) do not affect our results.

### Titration of world model accuracy

In Fig. S3 world model accuracy was changed as follows. All input state sequences were generated from the true 3-regular graph. When world model inaccuracy was 0, the world model state graph matched this graph exactly. To increase inaccuracy, every outgoing connection had its downstream target randomly re-assigned with probability *p*_*swap*_, corresponding to the x-axis in Fig. S3.

### Probability of overlap of HD codes

Assuming **r**_*α*_ and **r**_*β*_ are random *K*-sparse binary encodings of two different states *α* and *β*, and *K ≪ N*, then number *n*_*shared*_ of overlapping elements is Poisson distributed:

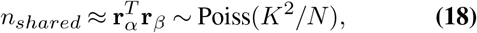

since for each 1 in **r**_*α*_ there is a probability of about *K/N* that there is a 1 at that location in **r**_*β*_, and there are *K* independent chances for this to occur. The expected overlap is

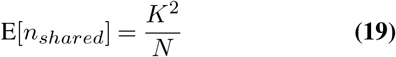

which is much less than *K* (the number of 1’s a sparse HD vector shares with itself) in the *N →∞* limit. Thus, 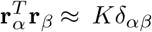.

### Capacity

How many states *L*^*\**^ can be stored in the readout layer? If we treat each state’s HD representation **r** as a pseudorandom *K*-sparse binary vector, our state-storage model is approximately (exactly in Fig. S2b, Fig. S2c) equivalent to a Bloom Filter, a well known distributed data structure for set membership queries (54), which has a false positive rate (equivalent to false familiarity detection) of

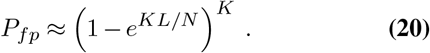

For a fixed false positive rate and activation density *K* this implies *L/N* must be constant, thus *L*^*\**^ *≥ 𝒪* (*N*).

### Encoding network parameters

Parameters are defined as follows. *M* : input (**u**) dimensionality; *N* : number of neurons; *Q*: connection probability between layers; *K*: activation density of final layer (corresponding to the *K*-WTA operation); *G*: gain between layers; *G*_*in*_: gain on input layer; *D*: number of layers. Values of these parameters for all results and figures are reported in Table S1.

### Familiarity detection and path retracing parameters

A threshold of *θ* = *K −* 1 was used in all simulations where the path was retraced for recall. In pure familiarity detection analyses (Fig. S2), the threshold was varied in the computation of the area-under-receiver-operating-characteristic (AU-ROC) metric.

## Acknowledgments

We would like to thank the many people with whom we had fruitful discussions about the development of this work, including: Kyle Aitken, Isabel Berwian, Tankut Can, Jonathan Cohen, Kayvon Daie, Francesco Fumarola, Michael Kahana, Misha Katkov, Kamesh Krishnamurthy, Nicolas Lenner, Agostina Palmigiano, Jonathan Pillow, Marton Rosza, Anna Schapiro, Dhairyya Singh, Julia Steinberg, Lee Susman, Misha Tsodyks; and other members of the Kahana, Schapiro, and Pillow groups. We also thank Matthew Rosenberg for a close reading of this manuscript, and Matthew Farrell and Alison Weber for reading an earlier version.

This work was supported in part through the Swartz Foundation, and by the National Science Foundation through the Center for the Physics of Biological Function (PHY-1734030). It was originally conceived at the espresso machine in the University of Washington Computational Neuroscience Center.

