## Supplementary Materials for "A non-Hebbian code for episodic memory"

| Simul. | $M$ | $N$ | $Q$ | $K$ | $G$ | $G_{in}$ | $D$ |
|---|---|---|---|---|---|---|---|
| Fig. 2d | 50 | 500 | .1 | 10 | 20 | 10 | 3 |
| Fig. 2d (L) | 50 | - | .1 | 30 | 20 | 10 | 3 |
| Fig. 2e (R) | 50 | 500 | .1 | - | 20 | 10 | 3 |
| Fig. 2h | 50 | - | .1 | 10 | 10 | 15 | 4 |
| Fig. 3e (L) | 50 | 1000 | .1 | 10 | 20 | 10 | 4 |
| Fig. 3e (R) | 50 | 1000 | .1 | - | 20 | 10 | 4 |
| Fig. 3g | 50 | 2000 | .1 | 10 | 10 | 10 | 10 |
| Fig. 3h | 50 | - | .1 | 10 | 10 | 20 | 10 |
| Fig. 3i | 50 | 2000 | .1 | 10 | 10 | 10 | 6 |
| Fig. 4b | 50 | 5000 | .1 | 10 | 10 | 20 | 10 |
| Fig. 4d | 50 | 3000 | .1 | 10 | 10 | 20 | 10 |
| Fig. 4h | 50 | 4000 | .1 | 10 | 10 | 20 | 10 |
| Fig. 5 | 10 | 4000 | .1 | 10 | 10 | 10 | 10 |
| Fig. S1a | 50 | 2000 | .1 | 10 | 10 | 10 | - |
| Fig. S1b | 50 | 2000 | .1 | 10 | - | 10 | 4 |
| Fig. S2b (T/B) | 50 | -/500 | .1 | 30/- | 20 | 10 | 3 |
| Fig. S2c (T/B) | 50 | -/500 | .1 | 30/- | 20 | 10 | 3 |
| Fig. S2d (T/B) | 50 | -/4000 | .1 | 30/- | 20 | 10 | 3 |
| Fig. S2e (T/B) | 50 | -/500 | .1 | 30/- | 20 | 10 | 3 |
| Fig. S3a | 50 | - | .1 | 100 | 20 | 10 | 4 |
| Fig. S3b | 50 | 1000 | .1 | 50 | 20 | 10 | 4 |
| Figs. S3c to S3d | 50 | 2000 | .1 | 10 | 10 | 10 | 6 |
| Fig. S4 | 10 | 4000 | .1 | 10 | 10 | 10 | 10 |

**Table S1.** Parameters values across all simulations and results for each figure. Dashes indicate parameter was varied during analysis.

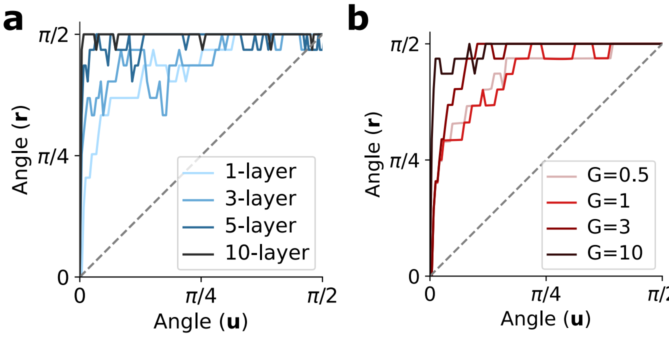


**Fig. S1. Locality-sensitivity of pattern separation by random feed-forward networks.** a. Angle between pair of output vectors $\mathbf{r}$ vs angle between pair of input vectors $\mathbf{u}$ as a function of number of layers in the network. b. As in A but as a function of gain $G$, for a 4-layer network.

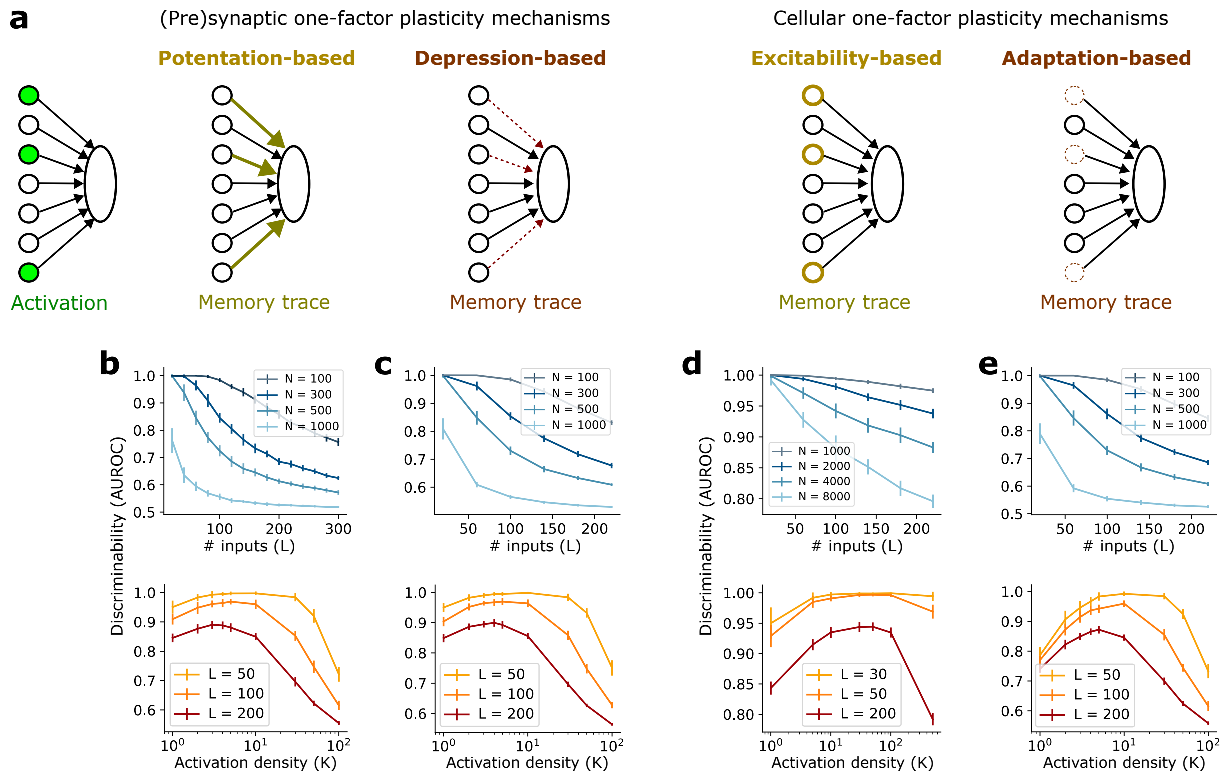


**Fig. S2. Four varieties of one-factor plasticity mechanisms with similar function.** a. Schematic of synaptic and cellular one-factor mechanisms. b. Familiarity detection through synaptic potentiation with bounded weights ($0 \leq w_i \leq 1$). Discriminability is assessed via area under the receiver operating characteristic curve. c. Familiarity detection through synaptic depression. Thin dashed lines indicated depressed synapses. d. Familiarity detection through instrinsic plasticity of excitability. Thick yellow outlines indicate excitability increases. e. Familiarity detection through cellular adaptation. Thin dashed outlines indicated adaptation (equivalently decreased excitability).

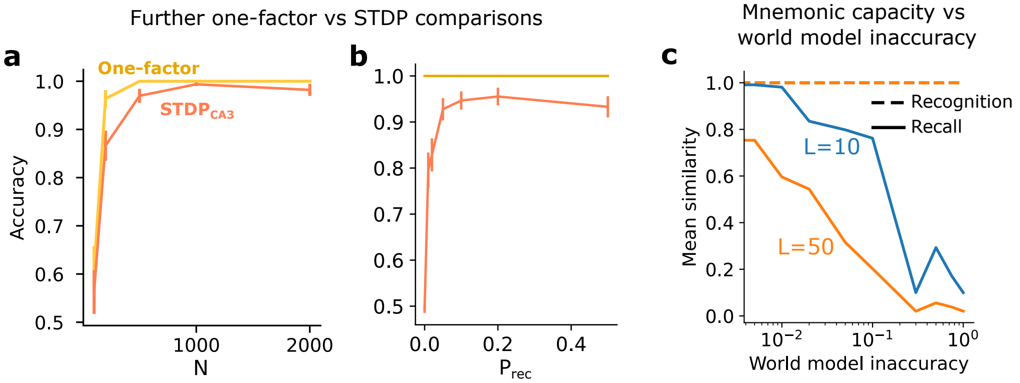


**Fig. S3. Further STDP comparisons and mnemonic capacity vs world model accuracy.** a. As in Fig. 2e except varying the network size. b. As in Fig. 2e except varying the recurrent connection probability. Fixed parameters for simulations in a, b are given in Table S1. c. Mean similarity between stored and recalled trajectories as a function of world model accuracy for a 3-regular directed graph with 5000 states. Dashed line shows recognition accuracy (correct familiarity detection). World model inaccuracy is defined as the probability of an outgoing connection in the state space having its target state randomly reassigned.

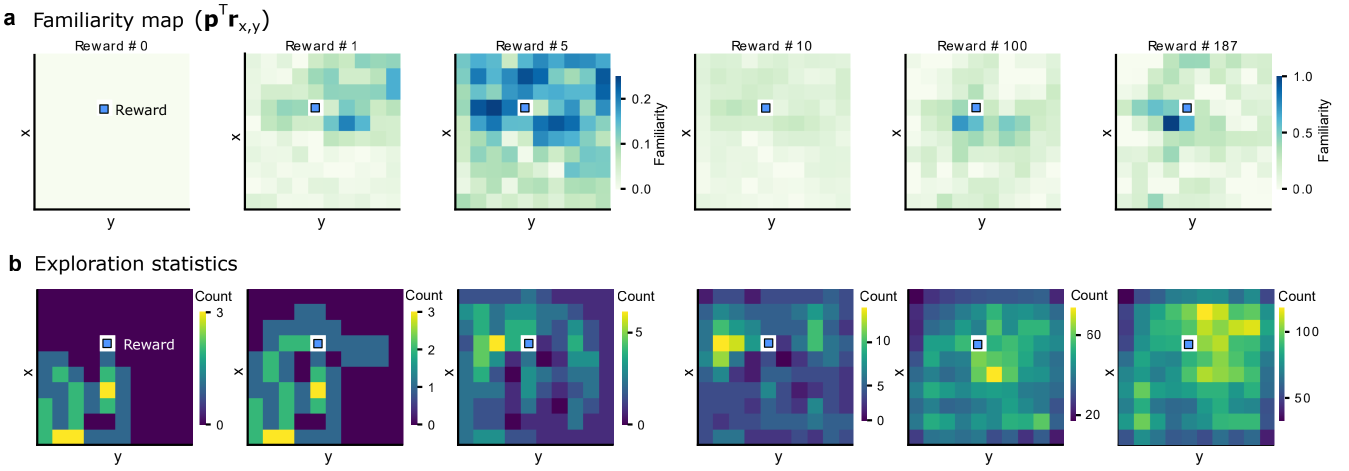


**Fig. S4. Off-policy learning with path vectors.** a) Familiarity maps during learning for off-policy behavior. Same as Fig. 5b for on-policy behavior. For each location in the environment a familiarity can be decoded through the learned path vector familiarity decoder. As learning proceeds, locations closer to the reward become increasingly more familiar. b) Statistics of explored locations during learning. This is one of the leading factors that drives learning.
